# Conservation and discovery of regulatory motifs across oomycetes through comparative genomic analysis

**DOI:** 10.1101/2025.02.18.638864

**Authors:** Sakshi Bharti, Marco Thines

**Affiliations:** Department of Biological Sciences, Institute of Ecology, Evolution and Diversity, Goethe University, Max-von-Laue-Str. 9, 60323 Frankfurt (Main), Germany; Senckenberg Biodiversity and Climate Research Centre (SBiK-F), Senckenberg Gesellschaft für Naturforschung, Senckenberganlage 25, 60325 Frankfurt (Main), Germany; Center for Integrative Fungal Research (IPF), Georg-Voigt-Str. 14-16, 60325 Frankfurt (Main), Germany; LOEWE Centre for Translational Biodiversity Genomics, Georg-Voigt-Str. 14-16,60325 Frankfurt am Main, Germany

**Keywords:** Oomycetes, transcription factors, regulatory elements, computational analysis, motif plasticity, pathogenesis-related mechanisms

## Abstract

Promoter sequences contain specific transcription factor (TFs) binding sites that regulate gene expression. While the conservation of TFs in pathogen development and infection among oomycetes is known, little is understood about TFs bind to conserved promoter regions across species. This study employs a robust comparative computational genomics approach to identify the TFs binding to orthologous DNA motifs in oomycetes. By integrating high-confidence TF binding site (TFBS) profiles, *in-silico* motif discovery, sequence conservation analysis and protein sequence similarity searches, the study revealed conserved regulatory mechanisms in oomycetes. The multi-layered computational framework identified two major TF classes in oomycetes: Cys2-His2 (C2H2) zinc finger proteins and winged helix repressor proteins, binding to orthologous motifs regulating gene clusters involved in epigenetic regulation, effectors, intracellular trafficking, host cell wall degrading enzymes, RNA processing and cytoskeletal organization. Structural comparisons indicate high sequence similarity between oomycete TFs and well-characterized eukaryotic TFs, supporting the predictive power of the computational approach. Moreover, motif plasticity analysis across developmental phases revealed conserved and phase-specific motifs emphasizing dynamic transcriptional regulation during infection and colonization. The presence of highly conserved motifs across multiple oomycete species suggests strong evolutionary selection pressure on key regulatory elements. The results provide a computational foundation for future experimental validation, guiding functional characterization of transcriptional regulation in oomycetes. This study highlights the potential of *in-silico* TFBS discovery for understanding gene regulation, paving the way for targeted experimental approaches such as ChIP-seq or electrophoretic mobility shift assays (EMSA).

## Introduction

Oomycetes, a group of filamentous eukaryotic pathogens, cause devastating plant diseases namely downy mildew, root rot and late blight (1,2) While morphologically resembling fungi, oomycetes are phylogenetically more closely related to brown algae and diatoms (1–4). Some of devastating plant pathogenic oomycetes, such as *Plasmopara halstedii* and *Phytophthora* spp., exhibit high genomic plasticity, which enable them to adapt rapidly to host environments (5–8). Despite their agricultural significance, the transcriptional regulatory mechanisms in oomycetes remain largely unexplored.

Transcription factors (TFs), are DNA-binding proteins that control transcription, by binding to specific DNA sequence sites within promoter region (9,10). Therefore, TF-DNA interactions serve as key regulators of gene expression across broad spectrum of life, ranging from bacteria to eukaryotes (Yang, 1998; Abril et al. 2020; Brázda et al. 2021). Promoter regions in eukaryotes typically contain conserved elements such as Initiator (Inr), BRE_d_ (Downstream TFIIB Recognition element), CAAT box, GC box, while motifs like TATA box, BRE_U_ (Upstream TFIIB Recognition element) show more variability (11,12). In oomycetes, the canonical TATA box is often replaced by Initiator-like sequence (INR) flanked by a novel FPR sequence and a downstream DPEP element (Bharti and Thines 2023; Bharti et al. 2023; McLeod et al. 2004; Pieterse et al. 1994; Roy et al. 2013b, a; Seidl et al., 2012). These non-coding regulatory sequences control the transcription of key genes, particularly those involved in pathogenicity, effector secretion and host adaptation.

Motif plasticity plays a pivotal role in regulatory evolution by driving adaptive changes in transcriptional networks through sequence divergence in promoter regions (13,14). In oomycetes, this plasticity contributes to the evolution of pathogenicity, as evidenced by major duplication events in *Phytophthora* (15,16). The ability of pathogenic oomycetes to adapt and evolve is strongly influenced by motif plasticity, which facilitates variation in pathogenicity causing genes, thereby enhancing host infection and immune evasion. For instance, structural and functional flexibility in key motifs, including RxLR, dEER, and LxLFLAK, plays a crucial role in effector translocation, subcellular localization and host manipulation, ultimately shaping pathogen-host interactions (17,18). Well-studied TF families include homeodomain, zinc finger, Myb, leucine zipper, helix-loop-helix, heat shock factors (HSFs) and forkhead box proteins (19). For instance, HSFs bind TTC repeats across eukaryotes, including plants, human and oomycetes (20). This large-scale analysis identified and characterized TF binding specificities belonging to 14 TF families. Similarly, Cys2-His2 zinc finger proteins are highly conserved across plants, animals and fungi, with their DNA-binding domains maintaining strong structural similarity (21–26). This high level of sequence and structure conservation reflects the fundamental role of TFs in gene regulation.

Despite advances in Next Generation Sequencing (NGS) for *Peronospora*, *Phytophthora* and *Pythium* species, accurately predicting TF-DNA interactions from short-read sequencing data remain challenging (27,28). Regulatory regions, such as promoter sequences evolve more rapidly and leading to species specific variations in TF binding (29). Due to lack of more comprehensive chromosome-level genomes and faster evolutionary changes in promoter regions, comparative genomics can identify the conserved non-coding promoter sequences (30–35). Furthermore, comparative analysis assigns functions to un-annotated genes across multiple genomes and identify functionally important low-affinity binding sites (31,36). In the absence of experimental ChIP-seq data, the computational approaches such as position weight matrices (PWMs), motif clustering and homology-based predictions also offer valuable alternatives (37–41). Cross-species motif conservation analysis, combined with TF ortholog mapping, enhances the reliability of computationally inferred TFBS predictions.

This study hypothesizes that conserved transcriptional regulatory mechanisms regulate gene expression in phylogenetically related oomycete species. Specifically, orthologous DNA motifs serve as binding sites for TFs that control pathogenicity-related genes. By integrating motif plasticity analysis, comparative genomics and functional annotation, the study aims to identify conserved TF-DNA binding motifs in oomycete pathogens. The study examines the regulatory roles of TFBS clusters in controlling gene expression across infection-related transcriptional phases. Furthermore, the evolutionary conservation of motif architecture between oomycetes and other eukaryotic lineages have been assessed.

## Materials and Methods

### a) Preparation of gene expression dataset, orthologous TFBS motifs and the TF dataset

The data for conserved transcription factor binding sites (TFBS) motif in co-regulated gene promoters of *Pl. halstedii* and related *Phytophthora* species were collected from Bharti et al. (2023) and Bharti and Thines (2023). This study extends the methods established in previous analysis to further investigate orthologous TF and binding site relationship (Fig 1a). The time-series transcriptome data to key asexual developmental phases, such as zoospore release (ZS), establishment of infection (IF), colonization (CO), induction of sporulation (SP), was obtained from *Pl. halstedii* datasets (30). Motif occurrence across transcriptional phases (ZS, IF, CO, SP) was determined based on predefined expression thresholds. A motif was considered present in a specific phase if its associated gene cluster exhibited expression above a log2 fold-change threshold of ≥1.5 relative to the previous time-point of the specific phase. The regulatory motif data, specifically, orthologous TFBS motifs were retrieved and analyzed using MEME Suite (v4.2.0) alongside custom python, R and shell scripts (available at: https://github.com/sakshianil/Transcriptional_regulation_oomycetes).

**Fig 1.**
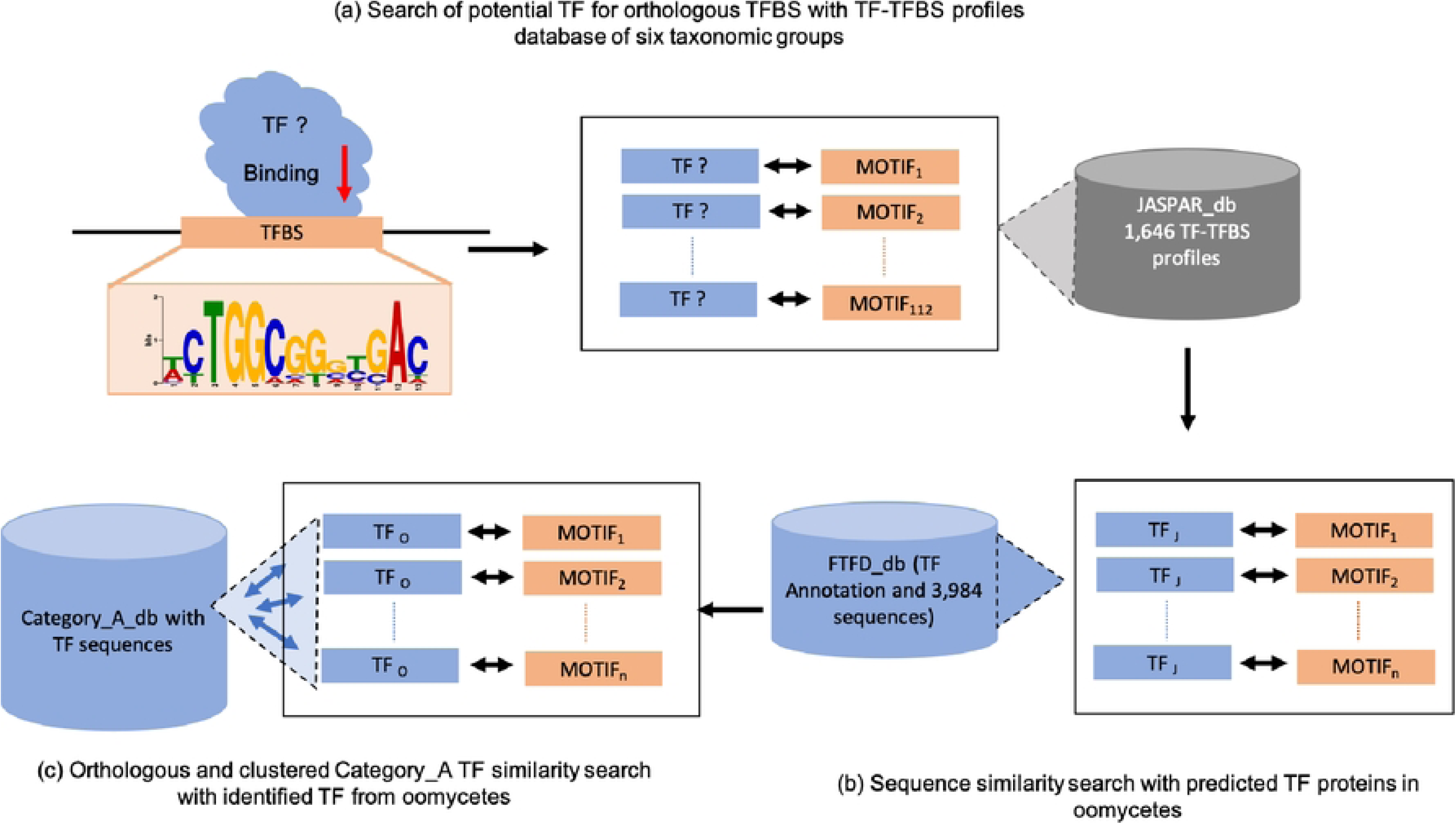
Workflow for transcription factor (TFs) identification and analysis of TF-binding sites (TFBS). The figure illustrates the data selection process, preparation of reference databases and identification of transcription factors linked to orthologous TF binding sites across *Plasmopara halstedii*, *Phytophthora sojae*, and *Phytophthora infestans*. (a) Screening of orthologous Category A motifs (TFBS), previously identified by Bharti and Thines (2023), against 1,646 eukaryotic TF profiles from the JASPAR CORE database to identify potential TF matches (E-value < 0.5). (b) Protein BLAST was used to compare statistically significant TF hits to 3,984 TF sequences from five oomycete species (*Hy. parasitica*, *Ph. sojae*, *Ph. capsici*, *Ph. ramorum* and *Ph. infestans*) in the Fungal Transcription Factor Database (FTFD), retrieving orthologous and homologous TFs (sequence similarity: 22–57%, E-value < 0.01). (c) Category A motifs, identified as predominantly TF-enriched gene clusters, were used to search the oomycete databases for associated TF proteins after no matches were found in JASPAR.

### b) Motif discovery and statistical enrichment analysis

To identify orthologous TFBS motifs, a two-tiered motif discovery strategy was employed: 1. Motif discovery with prior knowledge: A third-order Markov model was constructed as a background framework to improve motif identification in MEME. This statistical model helped differentiate true regulatory motifs from background sequence variation (30,31,42,43). 2. De novo motif discovery: The STREME algorithm (MEME Suite) was used for unbiased motif discovery in large gene clusters (≥50 members). Differential motif enrichment was assessed across developmental phases, pinpointing motifs conserved between species. This hybrid strategy ensured the identification of both conserved and plastic motifs, facilitating a comparative regulatory analysis.

### c) Preparation of reference TF-TFBS profiles from eukaryotic datasets

To ensure robust TF binding site annotation, experimentally validated TFBS profiles were sourced from Joint Annotated Source for the Prediction and Analysis of Regulatory regions (JASPAR) CORE databases (38,44). The selection criteria included: High-confidence, non-redundant TF binding profiles with experimental validation. Motif plasticity-based filtering using E-value thresholds at 0.1, 0.2, 0.5, and 0.05. There was inclusion of motifs across diverse taxonomic groups (*Vertebrata, Nematoda, Insecta, Plantae, Fungi, and Urochordata*). This dataset allowed for TF_J_-motif comparisons in phylogenetically distant species, offering a broader context for oomycete regulatory evolution utilized (Fig 1b).

### d) Preparation of oomycete-specific TF protein sequences

The translated and annotated transcription factor (TF_O_) protein sequences for five sequenced oomycete species — *Ph. infestans* (assembly ASM14294v1), *Ph. ramorum* (ASM14973v1), *Ph. sojae* (V3.0), *Hy. parasitica* (Phyt_para_P1569_V1) and *Ph. capsici* (LT1534 v11.0) — were retrieved from the Fungal Transcription Factor Database (FTFD) version 1.2 (Fig 1c; (19)). For TF proteins functional annotation, the verification methods used by FTFD were IEA (InterProScan, BLAST, PFAM or SMART).

### e) Comparative TF-TFBS analysis using eukaryotic and oomycetes reference databases

To evaluate the conservation of orthologous TF-motif interactions, 112 Category A motifs (31) were compared against JASPAR CORE motifs using Tomtom (MEME Suite v5.2.0) (38,42,44,45). These motifs were: previously identified in oomycetes and conserved across three species (*Pl. halstedii, Ph. infestans, Ph. sojae*). Derived from MEME (motifs with <50 gene members) and STREME (motifs with >50 gene members). Associated with gene clusters enriched in pathogenicity-related functions, including: effectors (e.g., RxLRs, CRNs), protease inhibitors, host cell wall-degrading enzymes (PCWDEs), RNA processing factors, intracellular trafficking genes, structural proteins, other TFs.

A log-likelihood scoring approach was applied to determine TF-motif homology. Additionally, the background letter frequencies were adjusted using motif-specific distributions. Motif comparisons were filtered using E-value < 0.5, ensuring statistically significant matches (45). Other filtering on parameters were E-value thresholds of 0.05, 0.5, 0.2, 0.1 and no small sample size correction. To expand motif-TF annotation, oomycete TF protein sequences were also compared with eukaryotic TFs from UniProt and compared with oomycete TFs from *Ph. infestans*, *Ph. ramorum, Ph. sojae*, *Hy. parasitica* and *Ph. capsici* using protein BLAST (BLASTp, E-value ≤ 0.01, word size = 7) (19,46–49).

### f) Direct sequence similarity analysis of oomycete-specific TFBS and TF proteins

For motifs not matching known eukaryotic TFBS profiles, direct sequence similarity searches were conducted using BLASTp against the oomycete TF database: BLASTp alignments were performed for TF-enriched motifs from *Pl. halstedii, Ph. sojae,* and *Ph. infestans*. Motif-TF associations were scored based on sequence identity, domain similarity and E-values (<3E-17). Pairwise alignments with the oomycete TF_O_ database were conducted using optimized BLAST parameters (46,47,49). This final step ensured that TF-motif relationships in oomycetes were evaluated even in the absence of direct eukaryotic orthologs.

## Results

### Retrieval of TF-TFBS profiles and protein sequence analysis of transcription factors across taxonomic groups

To investigate conserved transcription factor binding sites (TFBS) in oomycetes, 1,646 TF-TFBS profiles from the JASPAR CORE database were searched, identifying 38 unique eukaryotic transcription factors (TF_J_) across 47 motifs. These motifs were identified through MEME and STREME analysis, with statistically significant matches (E-value < 0.49; Appendix S1). The choice of E-value thresholds (0.05, 0.1, 0.2 and 0.5) follows previously established motif discovery frameworks, ensuring that biologically relevant, low-affinity binding motifs are not inadvertently excluded while maintaining statistical rigor (42,45,50). To improve specificity and exclude biologically relevant low-affinity binding sites, a more stringent threshold (E-value <0.05) was applied, refining the dataset to six motifs with 12 unique eukaryotic TF_J_ (Fig 2, 3). The identified motifs were significantly associated with biological functions including cell signaling, pre-mRNA splicing, RNA processing, PCWDEs, cellular metabolism, epigenetic regulation, effector activity, intracellular trafficking, protein degradation, and cytoskeletal and structural proteins (Appendix S1, S2). The predominant transcription factor families identified include basic leucine zippers (e.g., protein fosB, Creb5, opa), C2H2 zinc fingers, forkhead/winged helix factors (e.g., E2F6), homeodomain factors (e.g., MATALPHA2, HMRA1) and heat shock factors. The C2H2 zinc fingers and forkhead/winged helix factors TFs were found to be the most enriched classes, consistent with findings in fungal pathogens and higher eukaryotes, where they regulate stress responses, development, and pathogenicity (51,52).

**Fig 2.**
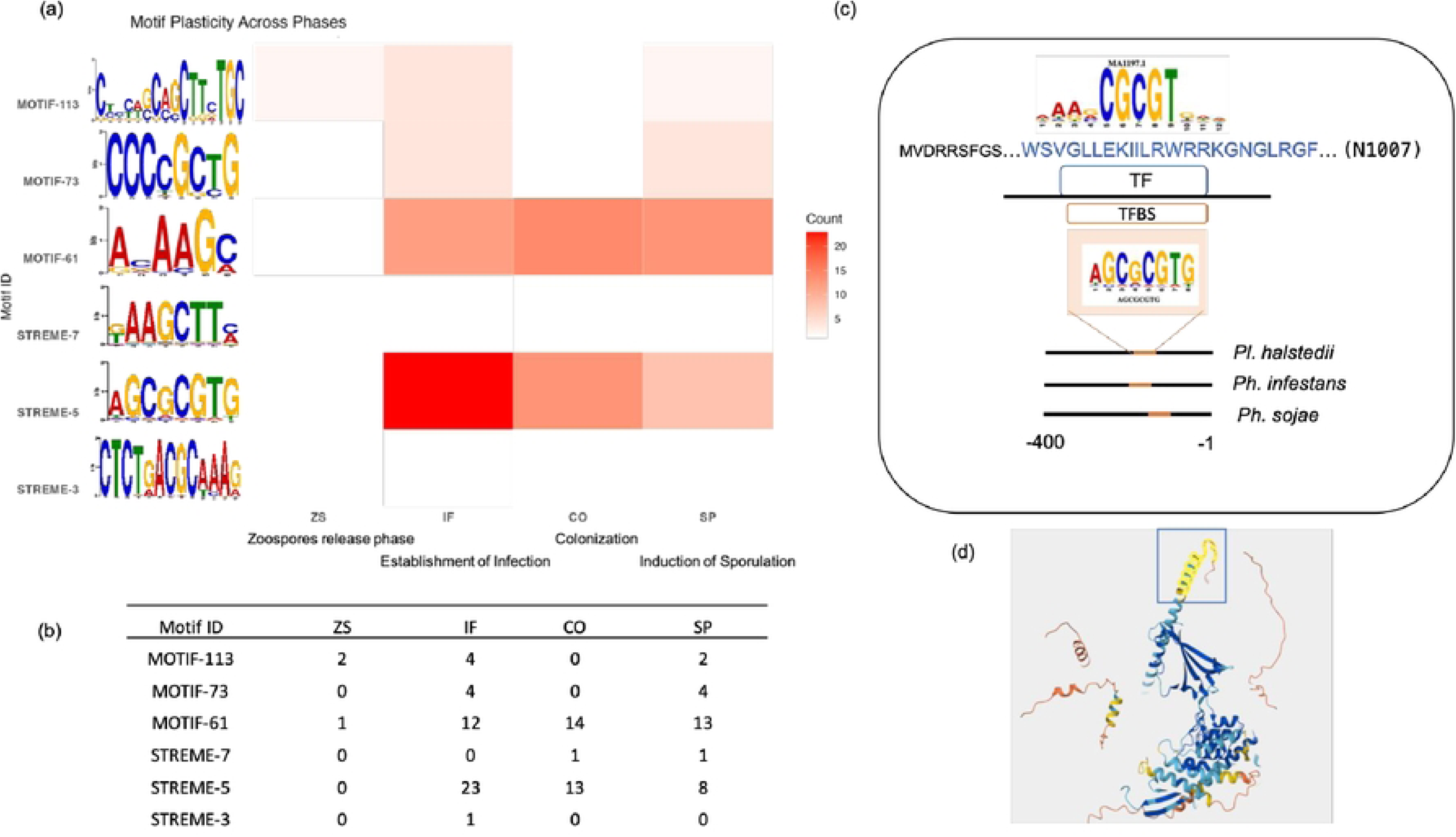
Motif plasticity and TF-binding Analysis. The figure illustrates the plasticity of motifs across transcriptional phases, their sequence logos, and the potential transcription factor (TF) binding interactions. The combination of heatmaps, sequence motifs, motif plasticity and structural visualization provides insights into the evolutionary conservation and regulatory importance of the identified motifs. (a) Heatmap of motif plasticity across asexual developmental Phases. The heatmap represents the occurrence of motifs across promoter regions of different transcriptional phases (ZS, IF, CO, SP). Color gradient: Higher intensity (red) indicates a greater number of occurrences of the motif in a specific phase. Each row corresponds to a specific motif-id or motif, labeled with its sequence logo, displaying nucleotide frequencies at each position. (b) A quantitative summary of motif conservation and plasticity across developmental stages of *Plasmopara halstedii*. Each row corresponds to a motif (E-value <0.05), labeled with its sequence logo, displaying nucleotide frequencies at each position. (c) TF-binding site mapping to eukaryotic DNA motif of JASPAR CORE database. A predicted transcription factor binding site (TFBS) derived from motif alignment (STREME-5; AGCGCGTG) is displayed. The consensus binding sequence of the TF is mapped to a highly conserved region of the motif in oomycetes. The identified DNA-binding residues within the TF sequence (blue text; Uniprot ID Q9FY74; 1007 amino acids; GCM domain factors) are aligned to the binding motif. (d) Structural representation of TF-DNA interaction. DNA motif is modeled within a predicted TF-DNA complex. The highlighted structural domain (yellow) corresponds to the region of the TF that directly interacts with the motif. The structure suggests potential protein-DNA interactions and binding affinity differences between motifs.

**Fig 3.**
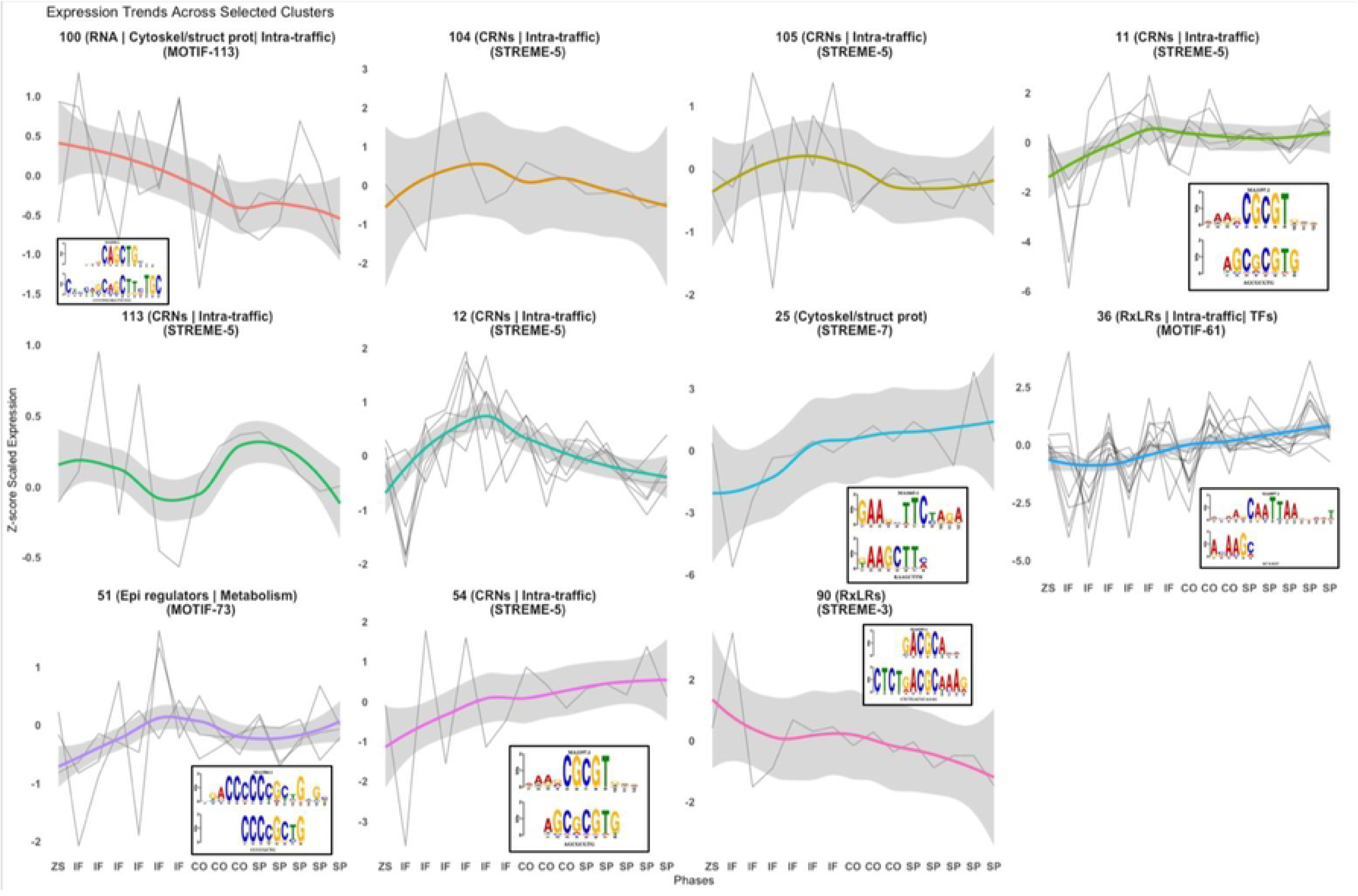
Motif-associated expression dynamics across clusters. This figure presents the expression trends of selected gene clusters across different transcriptional phases (ZS, IF, CO, SP) in *Plasmopara halstedii*, integrating motif occurrences with gene regulatory dynamics. Line plots representing cluster-specific gene expression trends: Each subplot represents a specific gene cluster, labeled with its functional category such as RxLRs (Effector Genes), CRNs (Pathogenicity-Related Transcriptional Regulators), Epigenetic Regulators, Intracellular Trafficking Components, Cytoskeletal and Structural Proteins. Abbreviations used: Intra traffic-Intracellular trafficking, RNA-RNA processing, Epi regulators-Epigenetic regulators, Cytoskel/struct prot: Cytoskeletal and structural proteins. Gray lines: Individual gene expression trajectories. Colored lines: Smoothed trend lines fitted using the LOESS method. Shaded regions: Confidence intervals indicating expression variability. X-Axis: Phases of Transcriptional Regulation. The x-axis follows a structured phase order: ZS → IF → CO → SP, reflecting biological transitions. The gene expression changes are mapped to four biological phases. ZS (Zoospore Release): Gene expression at 5 minutes (T5min) and 15 minutes (T15min). IF (Infection Establishment): Gene expression at time points T4h, T8h, T12h, T24h, T48h, and T72h. CO (Colonization Phase): Gene expression at time points T120h, T221h, and T288h. SP (Induction of Sporulation): Gene expression levels at T290h, T292h, T294h, T296h, and T296h_prim. Motif representations and their association with gene clusters (within black square boxes): Sequence logos display statistically significant motifs enriched in each cluster. Motifs are identified by their JASPAR-based sequence logos, with nucleotide heights indicating conservation strength. Each motif is labeled with its respective identifier (e.g., MOTIF-113, STREME-5, etc.), providing direct linkage to regulatory elements influencing gene expression.

A local database of 3,984 transcription factors from the Fungal Transcription Factor Database (FTFD) was further used for protein BLAST sequence similarity searches (19,46–49). This analysis identified 186 TF_O_ hits orthologous to 18 TF_J_ across eleven Category A motifs, with sequence similarity ranging from 22% to 57%, with significant matches (E-value < 0.01) (Appendix S2). These motifs regulate gene clusters associated with effectors, intracellular trafficking, cell signaling, PCWDEs, RNA processing, protein degradation, and cytoskeletal and structural proteins. Notably, while some TFBS motifs exhibited high conservation, other displayed species- or phase-specific plasticity, highlighting their potential role in transcriptional reprogramming.

### Motif stability and plasticity across transcriptional phases and structural insights into TF binding

To assess the temporal dynamics of motif activity, the distribution of motif across four transcriptional phases was analyzed: zoospore release (ZS), infection (IF), colonization (CO), and sporulation (SP) (Fig 2, 3, Table 1). Statistical significance for motif occurrence was determined using p-value, E-value and q-value (Appendix S1). IF and CO phases showed the highest motif retention, with several motifs (MOTIF-104, MOTIF-113, and STREME-5) being consistently detected in both, suggesting functional redundancy and co-regulation of genes involved in host colonization and metabolic adaptation. ZS phase exhibited the lowest motif occurrence, indicative of transient gene activation during early developmental stages. SP phase displayed the widest motif distribution, suggesting transcriptional reprogramming during sporulation. Among identified motifs, STREME-5 exhibited the highest occurrence in the IF phase (23 occurrences), suggesting a potential role in infection-specific gene regulation. Similarly, MOTIF-61 and MOTIF-79 were found across multiple phases, indicating their role in maintaining regulatory stability across different biological states. Structural analysis of STREME-5 further revealed alignment with a conserved transcription factor domain, emphasizing key amino acid residues involved in DNA binding (Fig 2c, 2d). The 3D protein structure analysis highlighted TF-DNA interactions, supporting the biological relevance of the identified motifs (Appendix S1, S2). This suggests that while conserved motifs may be essential for core regulatory processes, plastic motifs may contribute to environmental adaptation and host specificity.

**Table 1:** Phase-specific distribution and statistical significance of identified motifs. This table summarizes the positional occurrences, statistical significance, and phase-specific conservation of transcription factor binding motifs across four developmental phases: Zoospore Release (ZS), Infection (IF), Colonization (CO), and Sporulation (SP). The columns include Motif ID, Sequence, (ZS), (IF), (CO), (SP): Number of motif occurrences in each transcriptional phase, Start & End (ZS, IF, CO, SP; genomic coordinates of the motif within the respective phase): Genomic positions (in base pairs) where the motif was detected in each phase, “NA” indicates no occurrences in the respective phase, p-value (Statistical significance of motif enrichment, representing the probability of observing the motif by random chance), E-value (Expected frequency of motif occurrence by random chance in the dataset, lower values indicate higher confidence), q-value: False discovery rate (FDR)-adjusted significance level, correcting for multiple testing.

| Motif ID | Sequence | (ZS) | (IF) | (CO) | (SP) | Start<br>(ZS) | End<br>(ZS) | Start<br>(IF) | End<br>(IF) | Start<br>(CO) | End<br>(CO) | Start<br>(SP) | End<br>(SP) | p-value | E-value | q-value |
| --- | --- | --- | --- | --- | --- | --- | --- | --- | --- | --- | --- | --- | --- | --- | --- | --- |
| MOTIF-81 | CGCCWGCAGCKYC | 0 | 1 | 0 | 0 | NA | NA | 248 | 235 | NA | NA | NA | NA | 2.67E-04 | 4.39E-01 | 2.17E-01 |
| MOTIF-75 | YGTGGCGGGCAG | 0 | 6 | 5 | 4 | NA | NA | 222 | 210 | 205 | 193 | 198 | 186 | 2.74E-04 | 4.51E-01 | 4.35E-01 |
| MOTIF-56 | YCCCGGASGRCG | 0 | 13 | 0 | 7 | NA | NA | 250 | 238 | NA | NA | 304 | 292 | 1.89E-04 | 3.12E-01 | 2.34E-01 |
| MOTIF-133 | YCATGTAA | 0 | 8 | 6 | 0 | NA | NA | 283 | 275 | 282 | 274 | NA | NA | 1.23E-04 | 2.03E-01 | 1.63E-01 |
| MOTIF-85 | CGTM | 0 | 12 | 0 | 12 | NA | NA | 244 | 240 | NA | NA | 244 | 240 | 7.88E-05 | 1.30E-01 | 8.93E-02 |
| MOTIF-73 | CCCCGCTG | 0 | 4 | 0 | 4 | NA | NA | 301 | 293 | NA | NA | 355 | 347 | 3.84E-06 | 6.32E-03 | 3.01E-03 |
| MOTIF-101 | CCAGCCWSCC | 0 | 6 | 4 | 1 | NA | NA | 148 | 138 | 196 | 186 | 27 | 17 | 2.74E-04 | 4.51E-01 | 4.50E-01 |
| MOTIF-38 | AGAAKRYRATCAA<br>GG | 2 | 3 | 3 | 2 | 320 | 304 | 329 | 314 | 329 | 314 | 320 | 304 | 2.26E-04 | 3.71E-01 | 3.66E-01 |
| MOTIF-61 | ACAAGC | 1 | 12 | 14 | 13 | 125 | 119 | 217 | 211 | 211 | 205 | 213 | 207 | 2.48E-05 | 4.09E-02 | 4.09E-02 |
| MOTIF-124 | AAGCTACGGC | 3 | 6 | 0 | 0 | 363 | 353 | 292 | 282 | NA | NA | NA | NA | 2.36E-04 | 3.89E-01 | 2.91E-01 |
| MOTIF-46 | GGCAGCCCCGCTG<br>CA | 0 | 0 | 3 | 3 | NA | NA | NA | NA | 367 | 352 | 367 | 352 | 6.95E-05 | 1.14E-01 | 1.10E-01 |
| MOTIF-90 | AGCTGCTSCRTC | 0 | 6 | 1 | 4 | NA | NA | 204 | 192 | 253 | 241 | 214 | 202 | 1.45E-04 | 2.39E-01 | 2.38E-01 |
| MOTIF-65 | AGCTGCTGGW | 0 | 1 | 0 | 0 | NA | NA | 212 | 202 | NA | NA | NA | NA | 8.18E-05 | 1.35E-01 | 1.34E-01 |
| MOTIF-43 | CCCGCTGCA | 0 | 8 | 8 | 4 | NA | NA | 349 | 340 | 349 | 340 | 359 | 350 | 2.64E-04 | 4.35E-01 | 3.75E-01 |
| MOTIF-25 | AGCTACGGCAGCC<br>CC | 0 | 9 | 4 | 0 | NA | NA | 335 | 320 | 356 | 341 | NA | NA | 3.03E-04 | 4.99E-01 | 4.18E-01 |
| STREME-7 | KAAGCTTM | 0 | 0 | 1 | 1 | NA | NA | NA | NA | 91 | 84 | 91 | 84 | 3.02E-05 | 4.96E-02 | 1.79E-02 |
| STREME-5 | AGCGCGTG | 0 | 23 | 13 | 8 | NA | NA | 247 | 240 | 225 | 218 | 197 | 190 | 1.05E-05 | 1.72E-02 | 1.66E-02 |
| STREME-3 | CTCTGACGCAAAG | 0 | 1 | 0 | 0 | NA | NA | 116 | 104 | NA | NA | NA | NA | 1.44E-05 | 2.37E-02 | 2.34E-02 |

**Table 2.**
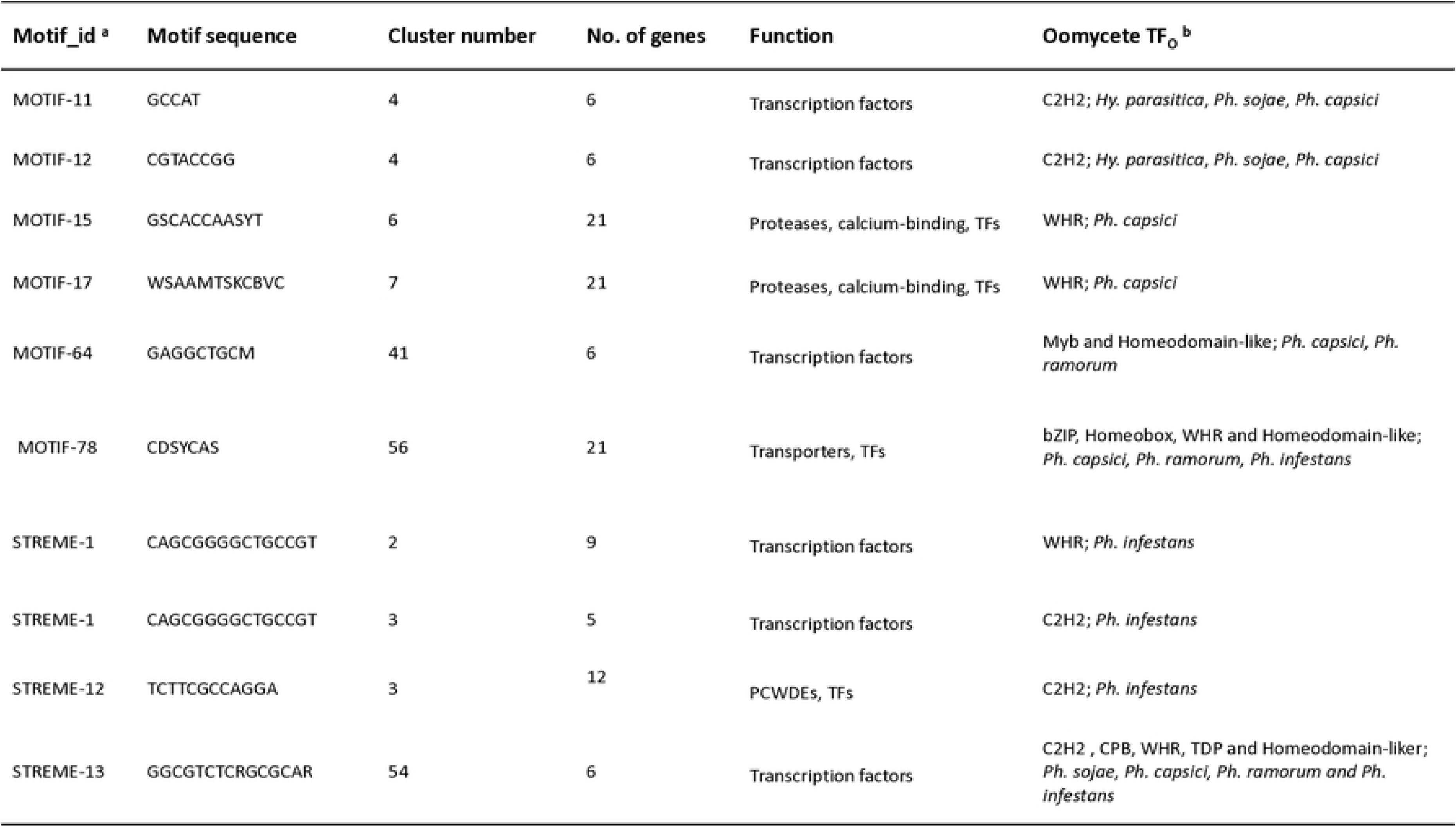
Potential and conserved transcription factor binding sites (TFBS) and transcription factors (TF_O_) retrieved from oomycete database (no matches were found in JASPAR database). The columns include motif id, sequence, cluster number, number of enriched genes, and their corresponding gene functions. Abbreviations: TFs= Transcription factors; Plant cell wall-degrading enzymes= PCWDEs; bZIP= Basic leucine zipper; TDP= Transcription factor E2F/dimerisation partner; WHR= Winged helix repressor DNA-binding; C2H2= C2H2 zinc finger; CPB= Centromere protein B, DNA-binding region; NAB= Nucleic acid-binding, OB-fold; HSF= Heat shock factor (HSF)-type, DNA-binding. ^a^ motifs from Bharti and Thines (2023). Represents the Category A motifs found to be upstream of the gene members of a cluster. ^b^ Conserved TFs in either or all of the oomycete species (*Hy. parasitica*, *Ph. sojae*, *Ph. capsici*, *Ph. ramorum and Ph. infestans*).

### Statistical significance of motif plasticity across phases

To assess the functional significance of motif variation across transcriptional phases, statistical thresholds were applied: Motifs with E-value < 0.05 were considered high-confidence regulatory elements. Motifs detected across three or more transcriptional phases were categorized as highly conserved. Phase-specific motifs were defined as those occurring in only one developmental phase. As shown in Fig 4, ZS-phase motifs exhibited the lowest p-values, indicating stronger statistical significance and higher conservation. In contrast: IF and CO phases displayed moderate motif significance (higher p-values), reflecting regulatory complexity during host colonization. SP phase showed the highest variability, suggesting extensive transcriptional rewiring during induction of sporulation.

**Fig 4.**
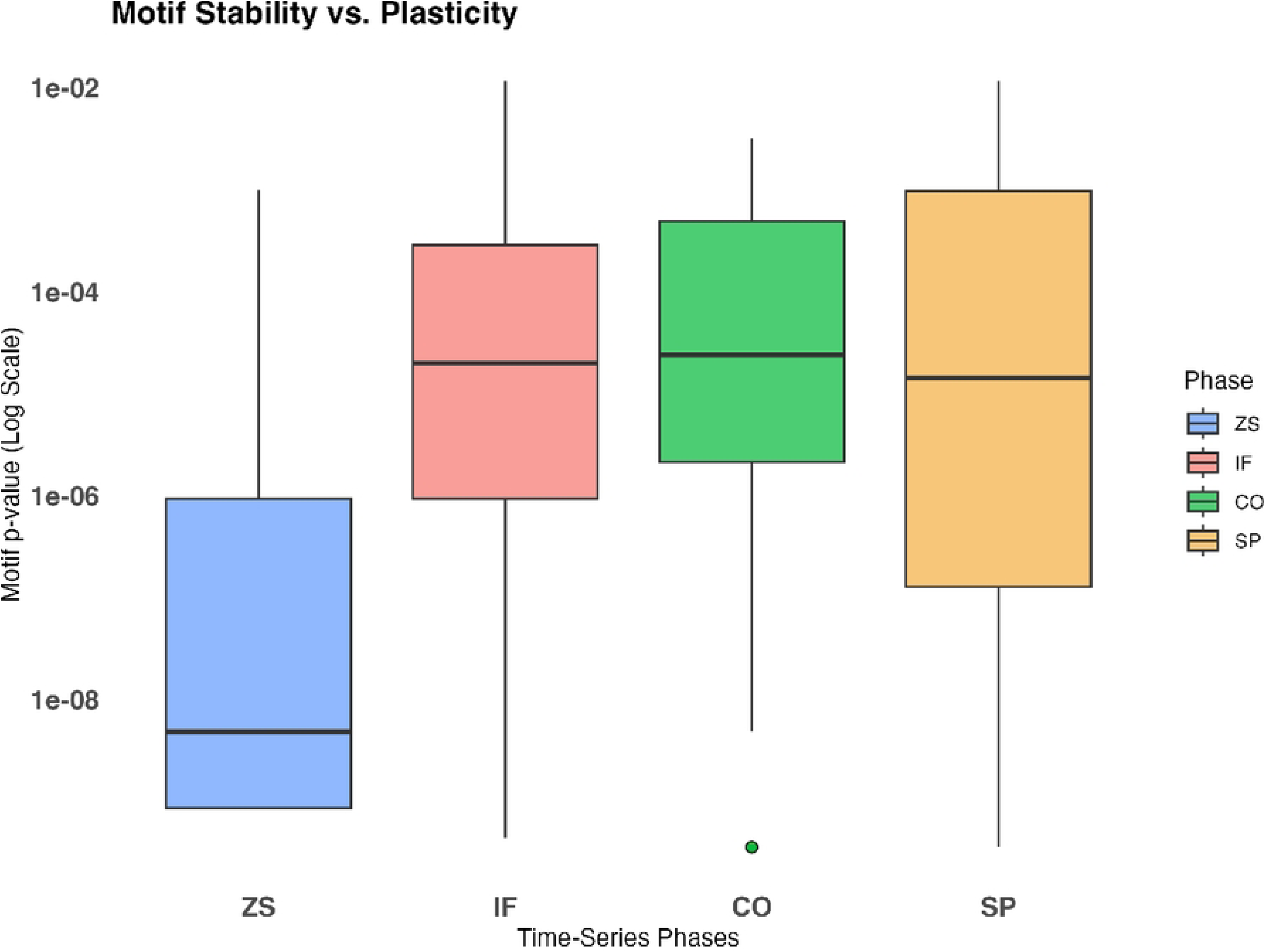
Motif stability vs. plasticity across phases. This boxplot visualizes the distribution of motif-associated p-values across four transcriptional phases: Zoospore Release (ZS), Infection Establishment (IF), Colonization (CO), and Sporulation (SP) in *Plasmopara halstedii*. X-Axis: Transcriptional Phases. Y-axis (log scale): Represents motif p-values on a log10-transformed scale, indicating statistical significance across phases. Each boxplot summarizes p-value distribution per transcriptional phase: Middle line (Median): Represents the median p-value. Outliers (dots): Represent highly significant motifs deviating from the main distribution. ZS (Zoospore Release - Blue): Initial phase of the oomycete life cycle, showing the lowest motif significance distribution. IF (Infection Establishment - Red): A transition phase with an increase in statistically significant motifs. CO (Colonization - Green): Higher median p-values suggest conserved motif activity in host adaptation. SP (Sporulation - Orange): A wider spread of p-values indicates motif-driven regulatory variability during sporulation.

### Functional redundancy and overlap in TFBS motifs across phases

Interestingly, certain motifs were identified across multiple phases, raising the question of functional redundancy among transcription factors. MOTIF-38, 61 and STREME-5 were observed in all four phases, suggesting their role as global regulatory elements. In contrast, MOTIF-81, 65 and STREME-3 displayed phase-specific enrichment, indicating specialized transcriptional functions (Appendix S1). This overlap suggests that TF families may share redundant roles, allowing for adaptive reprogramming in response to environmental signals. Similar redundancy has been observed in fungal pathogens, where zinc finger TFs control multiple transcriptional programs (53). This finding supports the hypothesis that pathogenic oomycetes utilize a complex regulatory network involving both conserved and plastic TFBS motifs.

### Sequence similarity search for Category A motifs for gene groups with no match with JASPAR

To investigate potential novel regulatory motifs, Category A motifs with no matches in JASPAR were further analyzed via BLASTp against the oomycete TF protein database. This search identified 37 transcription factors (TF_O_) across 19 loci corresponding to ten TF-enriched TFBS motifs. The primary TF families associated with these motifs include C2H2 zinc fingers and winged helix repressors, with an average sequence similarity of 35.7% and an E-value < 3E-17 (Appendix S3). The absence of these motifs in experimentally validated TF datasets suggests they may represent oomycete-specific regulatory elements. These findings provide compelling evidence for the existence of previously uncharacterized TFBS motifs that may play a role in oomycete pathogenicity.

## Discussion

### Transcription factor diversity and functional implications

Transcription factors (TFs) constitute a significant proportion (5–10%) of protein-coding genes in eukaryotic genomes (54). However, a time-series transcriptome analysis of *Plasmopara halstedii* revealed a lower percentage (1.6%) of TF-encoding genes (30). These TFs were clustered based on their differential expression patterns, uncovering potential regulatory networks controlling pathogenicity-related genes. A prior study by Bharti and Thines (2023), identified 46 out of 112 orthologous motifs regulating 25 gene clusters enriched with RxLR effectors, Crinklers (CRNs), proteases, protease inhibitors, transporters, intracellular trafficking components, calcium-binding proteins, and plant cell wall-degrading enzymes (PCWDEs). Notably, 18 orthologous DNA motifs were conserved across *Pl. halstedii*, *Ph. infestans*, and *Ph. sojae*, emphasizing their significance in TF-mediated transcriptional regulation, growth and pathogenicity.

### Predominance of C2H2 zinc finger and winged helix repressor proteins

Among the identified oomycete TFs, C2H2 zinc finger proteins represented the most abundant class (38.7%), followed by winged helix repressor DNA-binding proteins (28.5%). Heat shock factors (HSFs) and homeodomain-like proteins contributed smaller fractions (16.6% and 0.05%, respectively). The overrepresentation of C2H2 zinc finger proteins aligns with their known functions in eukaryotic transcriptional regulation, particularly in stress responses and developmental control. Their DNA-binding specificity and structural conservation across species further reinforce their functional importance.

### Motif occurrence and stability across developmental phases

Motif plasticity analysis revealed substantial variations in motif conservation across the four developmental phases: Zoospore Release (ZS), Infection (IF), Colonization (CO), and Sporulation (SP). Several motifs, such as MOTIF-61 (ACAAGC) and STREME-5 (AGCGCGTG), were consistently identified across multiple phases, suggesting their involvement in core regulatory functions (Fig 1b, Appendix Fig S1). In contrast, phase-specific motifs like MOTIF-79 (CGGCASCMCCRC) and STREME-7 (KAAGCTTM) were primarily identified in a single phase, indicating their potential role in stage-specific gene expression (Appendix S1).

Motif stability was highest in the IF and CO phases, with many motifs retained across both stages, suggesting functional redundancy or co-regulation of infection and colonization-related genes (Table 1). In contrast, the ZS phase exhibited the lowest motif occurrence, likely due to transient gene activation during early-stage development. The SP phase exhibited the widest motif distribution, with some motifs spanning multiple phases, indicative of transcriptional reprogramming during sporulation.

### Positional distribution of motifs in promoter regions

The spatial distribution of motifs across promoter regions further highlights their regulatory roles. Motif-104 (GYTACGGCAGCCCCG) and MOTIF-113 (CYYYWSCMGCTTCTGC) were predominantly found upstream of genes associated with infection, whereas MOTIF-72 (CTTCC) and MOTIF-82 (CRKACA) were enriched in colonization-related genes. STREME-3 (CTCTGACGCAAAG) and STREME-5 (AGCGCGTG) were identified near genes involved in sporulation, reinforcing their role in late-stage development. Interestingly, motif plasticity was observed in highly variable regions, such as those associated with effector genes. The co-occurrence of motifs in overlapping genomic positions, particularly in IF and CO phases (e.g., MOTIF-61 and MOTIF-79), suggests combinatorial regulation where multiple TFs coordinate gene expression. The presence of short, conserved motifs (e.g., MOTIF-81 and MOTIF-108) with low E-values suggests strong evolutionary selection for key regulatory elements.

### PCWDEs and CRNs enriched clusters and TF-mediated regulation

Regulatory motif enrichment in oomycetes revealed associations with PCWDEs and CRNs. In *Antirrhinum majus*, in vitro TF-DNA binding assays and DNA affinity purification sequencing (DAP-Seq) have validated the binding of bZIP proteins to hybrid C-box/G-box motifs (55–57). MOTIF-25 (cl 12; AGCTACGGCAGCCCC; putative C-box/G-box; 3/21 genes; 35 occurrences) identified in oomycetes shares similarity with the A. majus motif (GCCACGTCAGC) known to bind the basic leucine zipper 43 protein (BZIP43), which regulates glycoside hydrolases (PCWDEs) (Appendix S1). For oomycetes, bZIP proteins have been implicated in oxidative stress response (58,59). Additionally, the study identified MOTIF-85 (CGTM; cl 65; 24 genes; 28 occurrences), a regulatory motif linked to Crinkler proteins with a conserved LQLFLAK domain, bearing similarity to motifs with JASPAR IDs MA0840.1 and MA1131.1 (Appendix S1, S3). Additionally, motifs such as MOTIF-25 (C-box/G-box element) linked to PCWDEs and MOTIF-85 (CRN effector regulation) have been explicitly tied to their developmental role across phases (Table 1).

### Evolutionary conservation and functional significance of identified motifs

The motifs identified in this study were filtered using statistical stringency criteria to minimize false positives, ensuring reliable motif predictions. Several motifs identified in oomycetes resemble binding sites characterized in well-studied eukaryotes, suggesting functional conservation. For example: STREME-7 (KAAGCTTM) resembles *GAA* inverted repeats recognized by heat stress TFs HSFB2A, HSFC1, HSFA6b, and HSFB2B in *Arabidopsis thaliana* (60–62). However, the identified oomycete motif (gAAGCTTc) is a poor match to canonical *GAA* repeats, raising questions about its role in HSF binding (Appendix S1, S2). This divergence suggests lineage-specific adaptations in TF-DNA interactions. Interestingly, some motifs such as MOTIF-73 (CCCCGCTG) aligns with C2H2 zinc finger proteins and resembles *Drosophila* GACCCCCCGCTG, a motif linked to anterior gene expression during embryogenesis (63). Homology-based analyses indicate functional conservation between *Mus musculus* (ZIC1, ZIC3) and oomycete C2H2 TFs. MOTIF-133 (YCATGTAA) matches *HMRA1*, a yeast HOX-domain TF involved in cell-type regulation (64). This is suggesting that core transcriptional mechanisms might be conserved despite oomycetes’ unique evolutionary trajectory. Similarly, MOTIF-101 (CCAGCCWSCC) shares sequence similarity (35.5%) with ZNF460, a methylation-sensitive zinc finger TF in *Homo sapiens*, suggesting conservation of regulatory mechanisms across kingdoms (65). MOTIF-75 (YGTGGCGGGCAG), identified in oomycetes, aligns with E2F6, a mammalian transcriptional repressor involved in chromatin silencing. Additionally, winged helix repressors in oomycetes exhibit 39.6% sequence similarity with *E2F6*, indicating potential regulatory conservation (66,67).

### Implications for Oomycete Evolution and Adaptation

The observed motif plasticity aligns with previous findings indicating rapid evolution of non-coding regulatory regions in oomycetes (31). The presence of both conserved and plastic motifs across developmental phases supports the hypothesis that motif turnover contributes to adaptive gene regulation in response to host-pathogen interactions. Additionally, conserved cis-regulatory elements may serve as key targets for experimental validation and functional characterization. This study integrates motif plasticity with transcriptional regulation, providing insights into stage-specific and conserved TF binding sites in oomycetes. Additionally, comparative analysis with plant and fungal pathogens could reveal broader regulatory patterns underlying pathogenicity evolution.

### Conclusions

The identification of conserved DNA motifs and the associated transcription factor proteins across economically significant oomycete species offers valuable insights into the regulatory mechanisms that may be conserved throughout eukaryotic kingdoms. This study aimed to elucidate key components of the regulatory network, specifically targeting transcription factors (TFs) that bind to conserved transcription factor binding sites (TFBS) and regulate gene expression. The positional analysis highlights motif stability vs. plasticity across transcriptional phases. Some motifs maintain strict positional conservation, reinforcing their critical regulatory roles, while others display flexibility, suggesting adaptive binding to different regulatory elements. The major findings highlight the prominence of Cys2-His2 zinc finger proteins and winged helix repressor DNA-binding TF proteins associated with orthologous DNA motifs. These results suggest that such transcription factors may play crucial roles in developmental processes and pathogenesis across various oomycete species. While experimental validation using ChIP-seq or EMSA remains a future objective, the integration of evolutionary conservation, motif enrichment analysis, and expression correlation in this study provides strong computational evidence for regulatory significance. The insights gained may ultimately inform novel strategies for managing oomycete pathogens that present major challenges in agriculture.

## Statements & Declarations

### Ethics approval

Not applicable

### Consent for publication

The publisher Springer has the authors’ permission to publish their research findings.

## Data availability

All scripts and data generated and analyzed during this study are available in a public GitHub repository. The repository can be accessed at https://github.com/sakshianil/Transcriptional_regulation_oomycetes. This includes all code for computational analyses, as well as any additional data files relevant to the research. The repository is organized into subfolders for easy navigation, with detailed documentation provided in the ‘README.md’ file to assist users in understanding the structure and usage of the code. Any further inquiries about the data or methods used in this study can be directed to the corresponding author.

## Author Contributions

The computational experiment was conceived and designed by both authors. SB carried out the computational analysis, interpreted the results and wrote the manuscript. MT edited and approved the final version of the manuscript.

## Acknowledgements

We express our gratitude to the Thines laboratory group members for their valuable insights, discussions and helpful suggestions. We also thank LOEWE for providing the computational resources. We would like to extend special thanks to acknowledge Dr. Claus Weiland for his technical assistance in computational server access. We express gratitude to GRADE Language Service and Alison Davis for the proof-reading services in English language.

## Supplementary figures and files

**Appendix Fig S1: Motif plasticity across phases.**

This heatmap visualizes the distribution of motifs across different transcriptional phases in *Plasmopara halstedii*. The color intensity represents the motif count in each phase, highlighting motif enrichment and plasticity. Y-axis (Motif_ID): Lists the identified motifs (e.g., MOTIF-81, MOTIF-75, STREME-5, STREME-3). The motifs (E-value <0.5) are sorted based on their presence across phases. The transcriptional phases are on X-axis such as ZS (Zoospore Release): Motif occurrences during the early phase of pathogen development. IF (Infection Establishment): Motifs associated with host invasion and infection progression. CO (Colonization): Motifs contributing to pathogen growth within host tissues. SP (Sporulation): Motifs active during spore formation and pathogen dissemination. Color Intensity (Count Scale): Darker red shades indicate higher motif occurrences. Lighter shades represent motifs with lower counts or phase-specific regulation. White areas suggest motifs not detected in that phase.

**Appendix S1:** The file contains detailed information on Category A motifs conserved across *Plasmopara halstedii*, *Phytophthora sojae*, and *Phytophthora infestans*. The motifs were identified through MEME/STREME analysis conducted by Bharti and Thines (2023) and represent potential transcription factor binding sites (TFBS) associated with various gene clusters. The file includes details of Category A query motifs, corresponding target motifs from eukaryotic experimentally validated TF-binding site profiles, JASPAR IDs, UniProt identifiers, protein sequences, and genome coordinates for all motifs, along with supporting evidence such as PubMed references validating the motifs’ functional significance. The file contains motifs sorted based on significance thresholds (0.5, 0.2, 0.1 and 0.05), classification of highly stable vs. flexible motifs across Zoospore release (ZS), infection (IF), colonization (CO) and sporulation (SP) phases. Time-series gene expression profiles adapted from Bharti and Thines (2023). The data extracted from *Pl. halstedii* infection *Helianthus annuus*.

**Appendix S2:** The file provides a comprehensive overview of motif analysis results in *Plasmopara halstedii*, orthologs hemi-biotrophic species covering transcription factor (TF) associations, motif occurrences, positional distributions across different developmental phases and homologous motifs identified across various gene clusters in oomycete species. The file includes enriched gene groups, associated TFs, homologous motifs (compared to JASPAR and FTFD databases), UniProt identifiers, TF descriptions, and DNA-binding domains. Sequence alignments, alignment quality metrics and species details are presented.

**Appendix S3:** The file presents a sequence alignment analysis of transcription factors in oomycetes. The file includes alignments of motifs without hits in the JASPAR database, identifying transcription factors conserved across five oomycete species using protein BLAST. A comparative analysis of *Pl. halstedii* TF-proteins and their orthologs in related species using Clustal Omega, with conserved motifs identified through STREME/MEME, highlighting shared transcription factors.

